# Fast, safe and high-throughput Real-Time PCR protocol for molecular sex identification in *Salmo salar*, applicable to historic scale collections

**DOI:** 10.1101/2023.09.05.556299

**Authors:** Besnard Anne-Laure, Meslier Lisa, Jousseaume Thibaut, Nevoux Marie, Marchand Frédéric, Launey Sophie

## Abstract

Conservation and management of wild species needs a detailed understanding of their population structure and dynamics. For instance, there are increasing evidence for sex-specific life history trajectories, even in species with limited morphological sexual dimorphism. In these species, identifying the sex of individuals in some or all life stages is a methodological challenge, which requires the development of new tools. Building on recent findings in molecular biology, we developed a new protocol for sex identification in Atlantic salmon (*Salmo salar*), aiming at the following criteria: accurate sex identification, fast diagnostic, applicable to large sample size, applicable to degraded DNA extracted from e.g. fish scales, and safe for the operator by limiting health and safety hazard. The protocol relies on multiplex Real-Time PCR using a set of specific primers. We tested its accuracy on a test set of 90 DNA samples from individuals of known morphological sex, and demonstrated its applicability on a large DNA sample from 384 individuals of unknown sex.

## Introduction

Long-term individual-based monitoring of populations in the wild is a gold standard for research in ecology and evolutionary biology (Clutton-Block & Sheldon 2010). It generates phenotypic data that have key application for the conservation and management of species, e.g. by feeding stock assessment models. However, in weakly dimorphic species, sex is virtually an unobservable variable. As a result, models rarely account for sex-specific demographic processes, which lead to a lack of biological realism. In salmon, fishery management relies on the definition of conservation limits, often expressed in terms of a minimal egg deposition that will achieve long-term average maximum sustainable yield (ICES 2018). Information about sex-ratio is a prerequisite but very few empirical data is available, which prevent models to account for potential change in sex ratio over time or space (King et al. 2023). For instance, sex-ratio has not been updated since 1996 in the model for the management of French Atlantic salmon stock unit (Prévost et al. 1996).

Phenotypic sexing is not possible for most of salmon life as sexual dimorphism only takes place for a few months during the reproduction period. Rapid development in molecular sexing opens new avenues for ecology and management of wild populations. In particular, the analysis of historical collection of biological samples, such as fish scales collected as part of monitoring programs, offer the opportunity to investigate temporal change in sex-ratio. Indeed, DNA can be extracted from a fish scale, even reaching back dozens of years, but the DNA collected is often in small quantity and low quality, composed of small-size DNA fragments (A-L. Besnard, pers. com.). Genetic sex determination in salmonids conforms to a XX/XY male heterogametic system, but the sexual chromosomes cannot be discerned on a karyotype. Yano et al. (2013) identified a male-specific Y-chromosome gene, SdY, assumed to be the master sex-determining gene in salmonids. This finding allowed for the development of several molecular-based methods for identification of sex in Atlantic salmon *Salmo salar* and other salmonids. The rationale of these methods is to amplify part of the SdY gene, along with amplification of another marker, which acts as a control for DNA quality. If both markers are amplified, the individual is considered male, if only the control is amplified, the individual is considered female, if there is no amplification sex cannot be assigned.

Over recent years, several protocols have been proposed. For instance, Quéméré et al. (2014) developed SdY-specific primers to be included in a genotyping assay involving several microsatellite markers, but this approach is not cost/time effective when the analysis only targets sex determination. King & Stevens (2020) developed an assay amplifying a 700pb long fragment of the Y marker, along with a control designed in the fatty acid-binding proteins gene (Fabp6b) of 457pb. However, 700bp or 457pb fragments can be too long and thus unusable in the case of degraded DNA where fragments size may be shorter (e.g. DNA extracted from old scales). Their protocol relies on agarose gel runs, which is both time consuming and requires the processing of important quantities of DNA intercalant stain (SyBrsafe or BET), with associated health and safety concerns. Anglès d’Auriac & al. (2014) developed a method based on Real-Time PCR multiplex of the Y gene and a control marker designed in 18S control. Unfortunately, the control is not salmonid-specific as it amplifies all eukaryote DNA, including human and others species naturally present in environmental samples. The non-specificity of the control may return false positive females due cross-contamination problems. This limitation is exacerbated when working with degraded DNA in low concentration, such as found in old scale samples. Thus, available molecular sexing protocols still suffer some limitations for large scale application, in terms of practicality, cost, implementation time or accuracy on degraded DNA for instance.

In this paper, we present a fast and accurate sexing DNA protocol for Atlantic salmon, using multiplex RT PCR, based on Anglès d’Auriac & al. (2014) idea, but using primers for SdY marker and control gene that are both specific to salmonids and appropriate for short degraded DNA segments. This protocol allows high-throughput molecular sexing of large number of samples, while reducing the health and safety issues for laboratory personnel. The protocol relied on identifying the best combination of fragments length and composition for sexual and control gene adapted to our scales samples. The protocol was developed and its accuracy tested on samples from fish of known morphological sex. Finally, we present its application to a large number of samples from fish of unknown sex.

## Materials and methods

### Preliminary tests

The first step of our study is to identify the best combination of fragments length and composition for sexual and control gene. As we aim to apply our protocol to the degraded DNA that is generally extracted from scale samples, both fragments should be of small size (200pb or less). Also, to allow high-throughput application, we selected a multiplex approach for DNA amplifications using a Real-Time PCR Sybrgreen process. Given that sex identification is thus based on melting curve interpretation, the two fragments must have a distinct melting temperature, as a result of distinct amino-acid composition and length.

We tested several primers from the literature, or primers newly designed, to aiming at amplifying various sexual DNA fragment (table 1a) and various control fragments (table 1b). Each amplifying fragment have to be specific of salmonid species to avoid amplifying non-target DNA from other species. We used DNA extracted both from finclips and for scales, to assess the reliability of the protocol both on good quality (finclips) and degraded DNA.

**Table 1:**
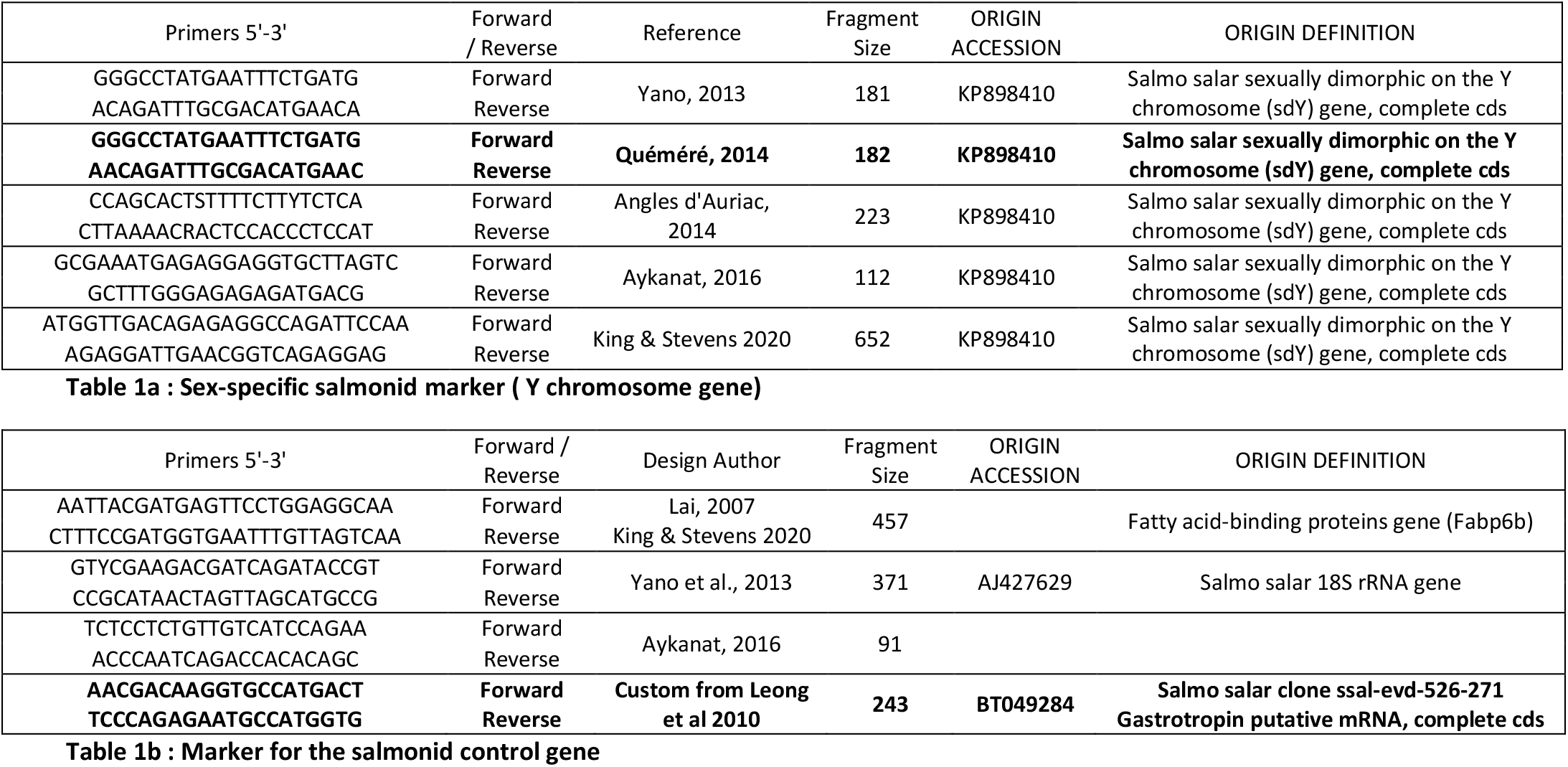
primer pairs for the sex-specific marker (1a) and the control marker (1b) tested in the preliminary test. In bold, the primer pairs chosen for the final protocol.

Some of the primers amplified well the focused fragment when run in isolation but returned a non-target amplifying (primer dimer) signal when run in multiplex. Some combinations of primers had melting temperatures that were too close to separate out the peaks from the sexual and from the control fragment. Others amplified well the focused fragments when DNA is extracted from finclips but did not perform well on the degraded DNA extracted from scales. From our tests, we selected a primer for a multiplex of sex marker Sdy of 182 pb (Quéméré et al., 2014) and designed a new specific control primer targeting a fragment of 243 pb within Atlantic salmon clone ssal-evd-526-271 Gastrotropin putative mRNA (FAB3) (Table 1).

### Final protocol for the best primer combination

DNA samples was extracted from different salmon tissues using the NucleoSpin® Tissue kit (Macherey-Nagel, no. 740741) according to the manufacturer instructions. Tissue was either finclip (1 mm^2^) fixed in alcohol, one scale for adult salmon or 2 to 3 scales for smolts. DNA concentration and quality was assessed on DeNovix® instrument, and DNA was systematically diluted 1:2 in DNase/Rnase free water to decrease inhibitors impact. Available final DNA concentration ranged from 0.78ng/μL to 50ng/μL.

A SybrGreen real-time PCR multiplex was performed in a total volume of 10μL. The mix contained 2μL of diluted DNA sample, final concentrations of 2X SsoADV Universal SYBR Green Supermix (BIORAD, n° 1725270) and 0.04μM of each primer. Thermocycling conditions were as follow: 95°C during 180 seconds, then 40 cycles of 95°C during 10 seconds and 60°C during 60 seconds. The analysis of melting curves was performed from 65°C to 95°C using increments of 0.5°C per second.

Validation of assays was based on the shape of amplification curves and melt-peak curves. We expect to observe amplification curves with exponential phase followed by a plateau and no more than two peaks positioned at the specific temperature on melt-peak curves. Any deviation from these criteria rendered the assay inconclusive. Sex identification is done from the observation of melt-peak curves. Males present two melt-peaks corresponding to the amplification of two fragments, sexual fragment Sdy and control fragment FAB3 at respectively 78,5 ± 0,5°C and 83,5 ± 0,5 °C. Females present only one fragment corresponding to the control fragment FAB3 (figure 1).

**Figure 1:**
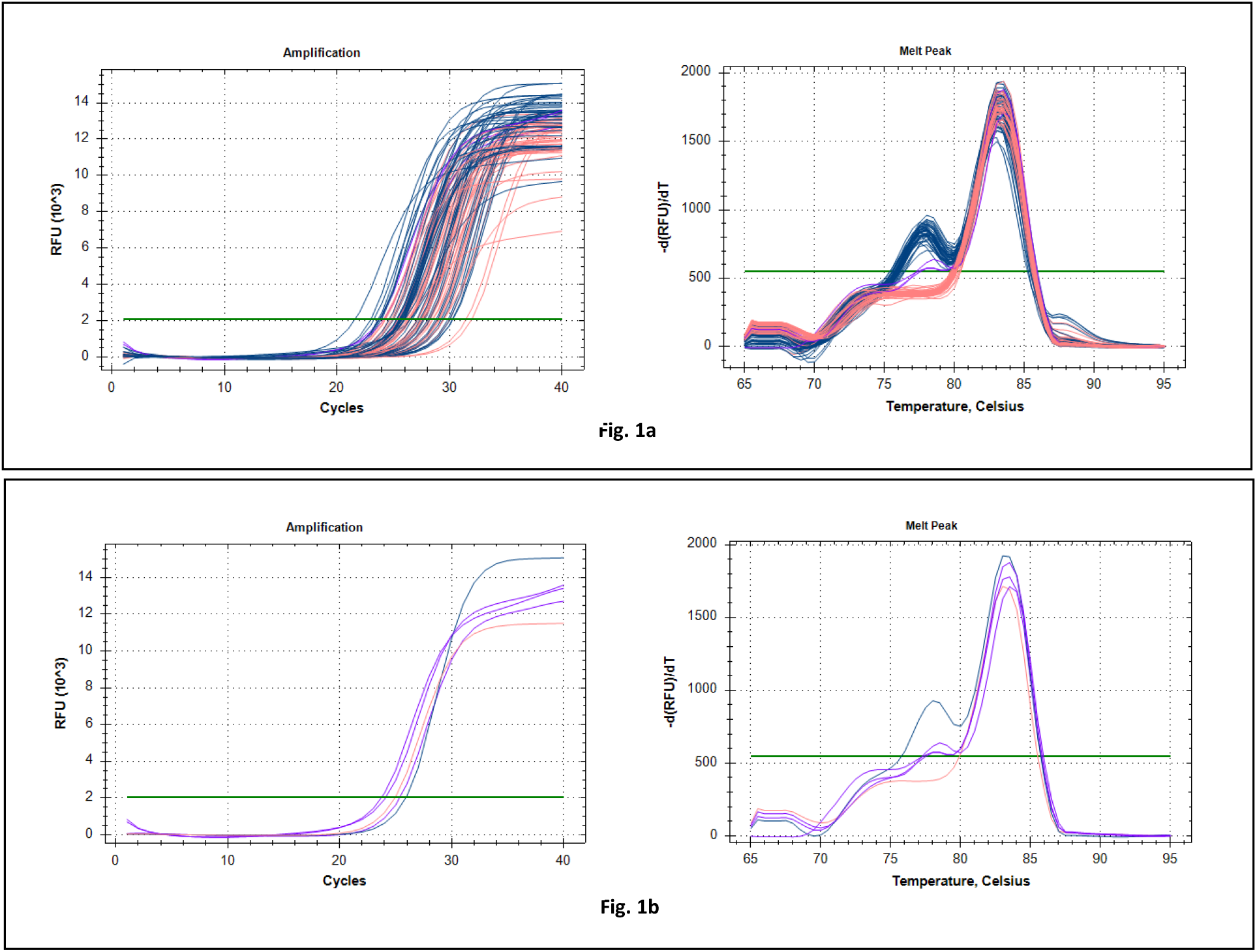
RT-PCR output curves for 90 known-sex individuals. 1a: Amplification curves (left) and Melt-peak curves (right) for all individuals. Males are coloured in blue, female in pink and non-characteristic profiles in purple. 1b: focus (in purple) on 3 non-characteristic amplification curve (left) and melt-peak curve (right) in comparison with characteristic male (blue) and female (pink) curves.

### Protocol validation

We assessed the accuracy of the protocol by testing its application on samples from Atlantic salmon of known phenotypic sex. DNA was obtained from finclips from 30 males and 30 females whose sex was formally identified from field observations (observation of sperm or eggs) during the 2018 breeding season on the Oir river (Normandy, France). Females were all large anadromous mature individuals caught on their return to freshwater after a marine sojourn. Ten males were small mature parrs that did not migrate at sea (precocious maturation) and 20 males were large anadromous mature individuals returning to freshwater after a marine sojourn. For 30 of these individuals (15 from each sex), we also extracted DNA from scales, to further test the accuracy of the analysis on degraded DNA. So, a total of 45 males and 45 females DNA samples were tested, corresponding to a total of 60 individuals. The genetic sex was determined from the above protocol, and compared with phenotypic sex after interpretation.

We then applied the protocol on the DNA extracted from 384 scale samples from wild Atlantic salmon of unknown sex. The scales were collected on individuals caught on the Bresle river trapping facility from 1990 to 2017 (table 3). The scales were randomly selected from the collection COLISA (Marchand et al. 2018) and encompassed different life stages (smolts and adults) and cohorts, in order to see if the protocol could be used on scales of various sizes and to assess the resolution rate of our protocol of molecular sexing in a large-scale analysis.

## Results

### Accuracy of the protocol on individuals of known sex

The selected amplified fragments are characterized by melt-peak temperature of 78.5 ± 0.5°C and 83.5 ± 0.5 °C, for the Sdy and the control fragments FAB3, respectively. Sex was identified without ambiguity on all 45 DNA samples from male Atlantic salmon (both finclips and scales), with the observation of exponential and plateau phase on amplification curves and two peaks on melt-peak curves (see Figure 1a).

Forty two out of the 45 female samples presented the expected female profile with an observation of exponential and plateau phase on amplification curves and a single control peak at 83.5 ± 0.5 °C on melt-peak curves (Figure 1a). Three samples (DNA from finclips) returned an ambiguous profile with an absence of the plateau phase and a small peak at the 78.5 ± 0.5°C target specific of the SdY melt-peak temperature (see Figure 1b and 1c). The protocol was run a second time on these three samples, which then returned the expected female profile.

### Upscaling the application of the protocol on a large sample of individuals of unknown sex

Out of the 384 DNA samples extracted from Atlantic salmon scales tested, 368 (95.8%) were unambiguously assigned to male or female with the first PCR assay (see table 2). The rate of sex resolution was lower for DNA samples extracted from old scales (90.3% for 1990-2002) than from recent scales (98.2% for 2005-2017). The rate of sex resolution was independent of the life stage (XX% in smolts and XX% in adults). Sixteen samples gave an ambiguous curve profile and sex could not be assigned in the first assay. We reported ambiguous profiles that were all characterised by the absence of a plateau, interpreted as a late amplification, an amplification of non-target fragment or the absence of amplification within the 40 cycles of the protocol (Figure 2). The assay was run a second time on those 16 samples and 14 of them then returned unambiguous results. After two runs, the rate of sex resolution reached 99.5%. The remaining 2 samples (0.5%) showed no amplification both times, which was attributed to faulty DNA.

**Table 2:**
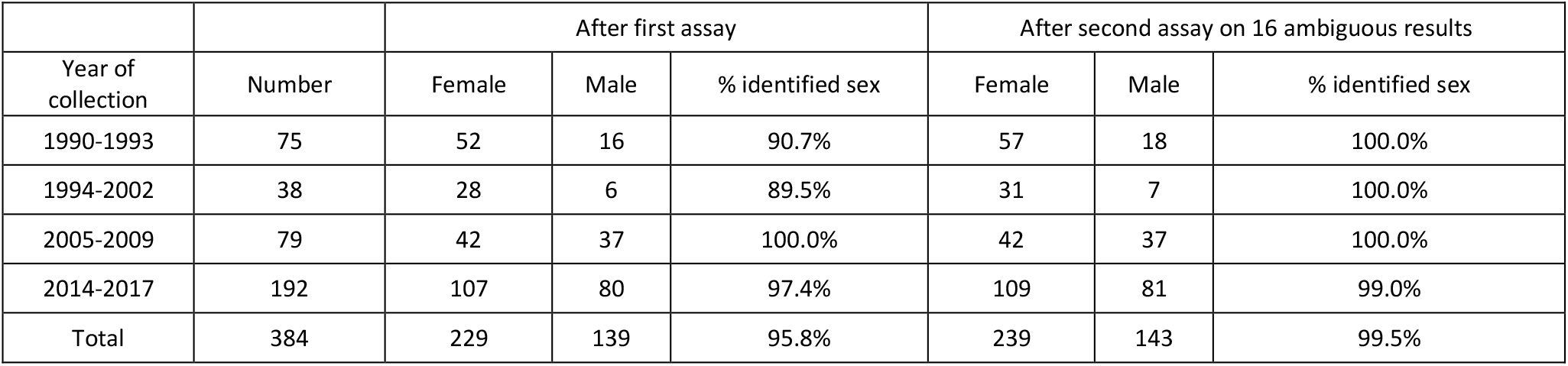
Sex identification results for 384 samples of unknown sex, collected between 1990 and 2017.

| Year of collection | Number | After first assay |  |  | After second assay on 16 ambiguous results |  |  |
| --- | --- | --- | --- | --- | --- | --- | --- |
|  |  | Female | Male | % identified sex | Female | Male | % identified sex |
| 1990-1993 | 75 | 52 | 16 | 90.7% | 57 | 18 | 100.0% |
| 1994-2002 | 38 | 28 | 6 | 89.5% | 31 | 7 | 100.0% |
| 2005-2009 | 79 | 42 | 37 | 100.0% | 42 | 37 | 100.0% |
| 2014-2017 | 192 | 107 | 80 | 97.4% | 109 | 81 | 99.0% |
| Total | 384 | 229 | 139 | 95.8% | 239 | 143 | 99.5% |

**Figure 2:**
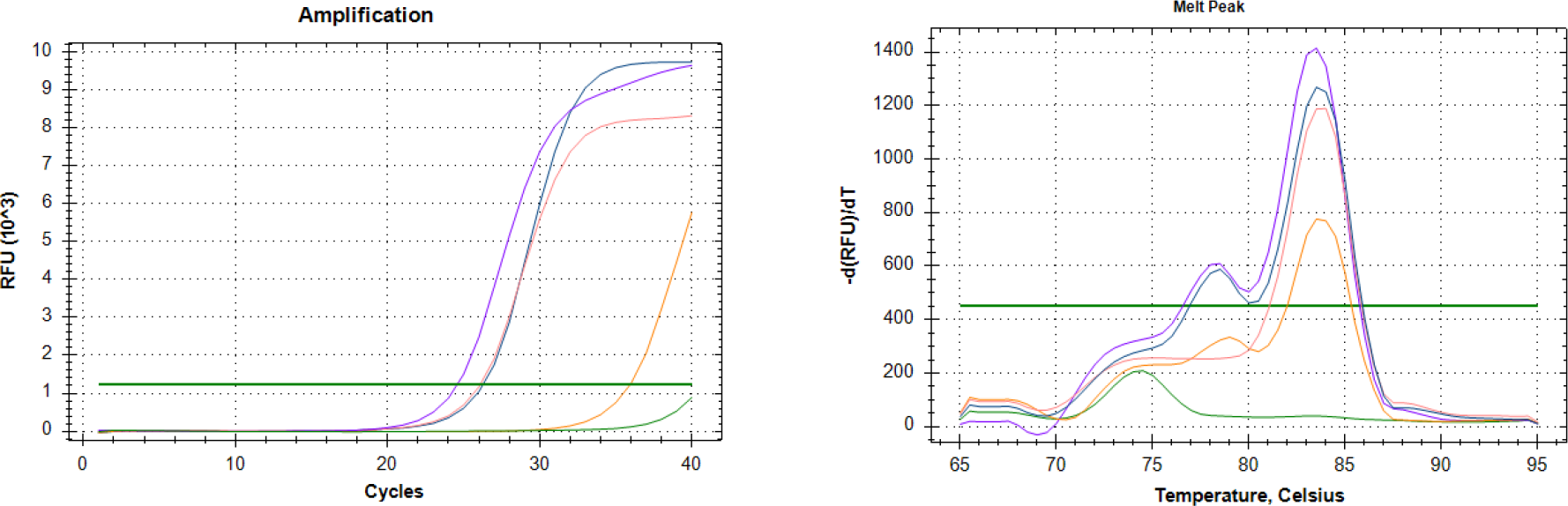
Example of various amplification curves (left) and Melt-peak curves (right) found during the sex determination on 384 samples from the scale collection. Characteristic female peaks in pink (229 found), characteristic male peaks in blue (139 found), non-specific profiles in purple, late profile in orange and no amplifying sample in green.

Within our random sample of Atlantic salmon caught at the trapping facility on the river Bresle, we detected XX% of females in smolts and YY% of females in adults.

## Discussion/conclusion

We present here an efficient molecular protocol to identify sex in Atlantic salmon from DNA extracted from historic scale collection. The protocol proved 100% accurate in sex identification for a subset of 90 samples from individuals of known phenotypic sex, using DNA extracted either from scales or finclips. When running the protocol on DNA extracted from scales that have been stored for more than 30 years, sex was unambiguously assigned in over 95% of the samples on the first assay. As a perspective, the reduced length of the selected primers makes our protocol robust for an application on DNA extracted from an extended variety of biological samples. Future investigations may allow reconsidering difficult or overlooked samples, such as mucus.

We obtained ambiguous amplification and/or melt-peak curve profile that depart from the typical male and female profile in 5% of the samples. When running the assay a second time on the same samples, uncertainty was lifted in most cases and the resolution rate of sex identification rises to 99.5%, even for scale samples dating back more than 20 years. The protocol is fast as it necessitates only one PCR run. The sex identification can be obtained for 96 individuals at a time in 1 to 1.5 days. It generates only limited hazardous waste, the only exception being the small quantity of SYBRGreen dye in the PCR mix.

Our protocol for sex identification in Atlantic salmon is designed based on the master sex-determining gene sdY, which is highly conserved in most salmonids (Yano et al. 2013). If the gene selected as a control is similarly conserved in these species, this protocol may be widely applied for sex identification in other salmonids too.

